# Polyisobutylene Micro/Nanoplastics Induce Multi-System Toxicity, Intestinal Barrier Dysfunction, and Neurodegeneration in *Drosophila melanogaster*

**DOI:** 10.64898/2026.09.18.752539

**Authors:** Darshini M. Basavaraju, Abass Toba Anifowoshe, Rounab Sarkar, Upendra Nongthomba

**Author notes:** Corresponding authors &. Shared First Author (Darshini M. Basavaraju, Abass Toba Anifowoshe).

## Abstract

Micro/nanoplastics (MNPs) are increasingly recognized as a persistent environmental contaminant of concern, yet polyisobutylene (PIB), an industrially significant polymer used as industrial sealants, adhesives, cable insulation, lubricant additives, and chewing gum bases, remains poorly studied relative to more commonly studied plastics such as polyethylene terephthalate (PET), polystyrene (PS), and polyethylene (PE). This study investigated the multisystem toxicological effects of chronic dietary exposure to PIB-MNPs in *Drosophila melanogaster* (*w¹¹¹⁸*). PIB-MNPs (> 5 μm) were synthesized via the solvent evaporation method and characterized using scanning electron microscopy (SEM) and Fourier Transform Infrared spectroscopy (FTIR), confirming successful synthesis. Flies were chronically exposed to PIB-MNPs (1, 2, or 3 mg) alongside a SDS control and a water control to 50 mg of yeast over 21 days, with toxicological outcomes assessed via survival analysis, climbing assays, fluorescence brain imaging, fertility/fecundity assays, the Smurf gut-permeability assay, Ellman’s AChE activity assay, and RT-qPCR analysis of *Stat92E* expression, a marker of inflammation. Chronic exposure to PIB-MNPs reduced survival, with females exhibiting markedly greater susceptibility than males. Exposure progressively impaired locomotor performance and induced brain vacuolization suggestive of neurodegeneration, peaking at day 15. AChE activity declined in a concentration-dependent manner by day 21. Female fecundity, egg-hatching rate, and ovarian egg-chamber maturation were significantly reduced, while male gonadal cyst cell numbers remained largely unaffected. PIB-MNPs exposure also induced a positive Smurf phenotype, indicating compromised intestinal barrier integrity and upregulated *Stat92E* expression, consistent with activation of a systemic stress response. Notably, several endpoints showed non-monotonic, concentration-independent trends, suggesting particle aggregation may influence effective bioavailability. Collectively, these findings establish PIB-MNPs as a multi-system toxicant in *Drosophila*.

**Graphical Abstract:** 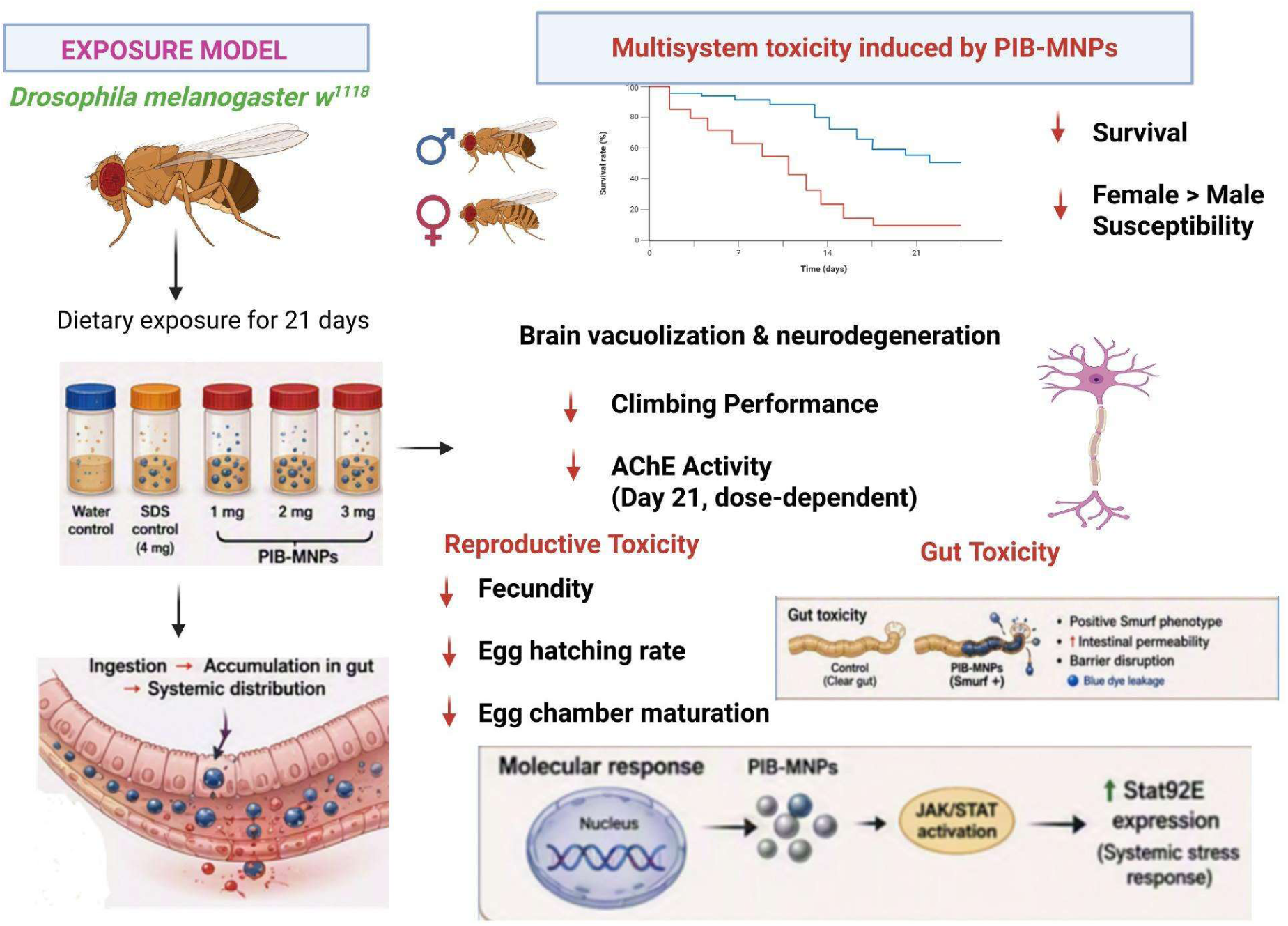

## Introduction

Polyisobutylene (PIB), also known as polyisobutene or butyl rubber when crosslinked, is produced through the cationic polymerization of isobutylene (2-methylpropene) at low temperatures, typically using Lewis’s acid catalysts such as AlCl₃ or BF₃ (Alves et al., 2021). The resulting polymer exhibits exceptionally low gas permeability, high electrical resistivity, good resistance to weathering and oxidation, and a broad viscosity range depending on molecular weight. These properties underpin PIB’s widespread application in automotive sealants, adhesives, chewing gum bases, lubricant additives (where it serves as a viscosity modifier), electrical cable insulation, and pharmaceutical tablet coatings (Bhatt et al., 2021).

Among the diverse array of polymers contributing to environmental micro/nano-plastic (MNPs) contamination, PIB-MNPs represent a chemically distinct and understudied class. Unlike the more frequently studied polystyrene (PS), polypropylene (PP), polyethylene (PE), or polyethylene terephthalate (PET) microplastics, PIB-derived MNPs have received comparatively limited toxicological characterization despite their worldwide applications and being detectable in environmental matrices. The hydrophobic, chemically stable nature of PIB-MNPs raises significant concerns regarding their bioaccumulation potential and capacity to induce oxidative stress, inflammatory responses, and genotoxic damage in biological systems.

Several *in vitro* studies on PIB toxicity have reported low or negligible adverse effects on cultured cells (Aarsaether, 1987). However, approximately two decades ago, Camphuysen et al. (1999) documented a major environmental incident in which PIB was implicated in the death of over 1,000 seabirds in the North Sea. In another earlier investigation, Iversen (1990) examined the tumor-promoting potential of various oils used to impregnate paper-insulated power cables, including PIB and its synthetic substitutes. Using male and female hr/hr Oslo strain mice, he applied 7,12-dimethylbenz(α)anthracene (DMBA) to hairless skin followed by PIB exposure. No evidence of tumor promotion was observed. Interestingly, the author concluded that 40% PIB oil may have reduced tumor incidence compared with DMBA alone or DMBA combined with 20% PIB oil. He further suggested that PIB increased mortality at higher dosages, although no supporting data were provided for this claim. Thus, direct *in vivo* evidence for the toxicity of PIB-MNPs remains scarce, but an important recent contribution comes from Anifowoshe et al. (2024), who synthesized pristine and fluorescently labelled PIB MPs in the size range of <2–10 μm using a solvent evaporation method and confirmed their structural stability by FTIR.

Exposing zebrafish (*Danio rerio*) larvae to a range of PIB-MPs concentrations, the authors reported delayed hatching, impaired swimming behaviour, elevated reactive oxygen species levels, abnormal spinal curvature, and reduced survival, alongside altered transcript levels of antioxidant-encoding genes including catalase (*cat*), superoxide dismutase (*sod 1 & 2),* and glutathione peroxidase (*gpx*). These findings constitute, to date, the most direct demonstration that PIB MPs are not merely persistent but actively toxic in a living vertebrate model, producing oxidative stress and developmental disruption through mechanisms broadly consistent with those documented for better-studied polymers such as PS and PE. The study nonetheless leaves open the neurological, reproductive, and gut-level consequences of PIB exposure, as well as the question of whether such damage can be mitigated by a co-administered protective agent, a gap that the present study, using an invertebrate model (*Drosophila melanogaster*), is designed to address.

*D. melanogaste*r occupies a uniquely powerful position in biological and toxicological research, combining extraordinary genetic tractability, short generation times, well-characterized neurobiology, and over a century of systematic experimental use (Prüßing et al., 2013). Approximately 75% of human disease-associated genes possess functional homologues in *Drosophila*, lending significant translational relevance to findings generated in this model (Atoki et al., 2025). Furthermore, the translucent larval and pupal stages, small body size, and rich repertoire of established behavioural and molecular assays render *D. melanogaster* exceptionally amenable to multi-level toxicological assessment. In recent years, *D. melanogaster* has gained widespread recognition as an eco-environmental model organism for assessing the toxicological impacts of microplastics (MPs) and nanoplastics (NPs). Dietary ingestion of plastic microparticles (such as polystyrene and polyethylene terephthalate) in *Drosophila* causes severe particle size-dependent intestinal damage, disrupts gut membrane integrity, and generates systemic oxidative stress (Matthews et al., 2021; Wang et al., 2024). Furthermore, because smaller nanoplastics can breach intestinal barriers and circulate systemically, exposure triggers widespread neuromuscular dysfunction, altered circadian locomotor activity, cardiac impairment, and sex-specific reductions in lifespan and reproductive output (Liu et al., 2022; Liang et al., 2022). *Drosophila* also provides a platform for investigating chemical co-exposure dynamics, demonstrating that microplastics can act as vectors that aggravate heavy metal toxicity and induce somatic epigenetic silencing (Sci. Total Environ., 2020).

To address the lack of systemic toxicity data for polyisobutylene, the present study evaluated the multi-system toxicological consequences of chronic dietary exposure to synthesized PIB-MNPs in *Drosophila melanogaster*. By combining physicochemical particle characterization (SEM and FTIR) with chronic 21-day exposure regimes, we systematically examined survival dynamics, locomotor coordination (negative geotaxis), whole-brain neurodegenerative vacuolization, acetylcholinesterase (AChE) enzymatic activity, intestinal barrier integrity (Smurf assay), male and female reproductive output, and transcriptional activation of the *Stat92E* stress-response signaling pathway.

## Materials and Methods

### *Drosophila* stock and culture maintenance

The *Drosophila melanogaster* white mutant line *w¹¹¹⁸* (Bloomington Stock (BS)# 5905) was used throughout this study. Stock populations were maintained in standard *Drosophila* culture bottles containing standard cornmeal-agar medium, composed of cornmeal, agar, yeast, sucrose, and propionic acid as an antifungal agent. All stock bottles and experimental glass vials were maintained in a temperature-controlled incubator set to 25°C with a relative humidity of 60-70% and a 12-hour light/dark photoperiod cycle (Gupta et al. 2025).

### Synthesis and Characterization of Polyisobutylene MNPs

PIB-MNPs were synthesized using the solvent evaporation method based on the procedure described by Anifowoshe et al. (2024), with slight modifications. Briefly, 500 mg of solid PIB (Santa Cruz Biotechnology, Catalog No.: SC-255434) was dissolved in 20 mL of chloroform (CHCl₃). Separately, 2 g of sodium dodecyl sulfate (SDS) was dissolved in 40 mL of double-distilled water (ddH₂O). Both solutions were mixed to form a biphasic mixture and sonicated for 30 minutes and oven-dried at 61.2°C. The resulting dry PIB-MNPs were collected and stored in a glass container for further characterization and analysis at room temperature.

The synthesized PIB-MNPs were subjected to comprehensive physicochemical characterization using a combination of analytical techniques to confirm their structural, morphological, and chemical properties. The surface morphology and structural features of the synthesized PIB-MNPs were examined using a Scanning Electron Microscope (SEM) (JEOL JSM-IT300), operated at an accelerating voltage of 15.0 kV and a working distance of 10.0 mm.

The chemical nature and functional group composition of the synthesized dry, powdered PIB-MNPs were further analyzed using Fourier Transform Infrared (FTIR) spectroscopy, performed in Attenuated Total Reflection (ATR) mode using a PerkinElmer Spectrum One system. The infrared transmittance spectra were recorded over a wavenumber range of 650 to 4000 cm⁻¹, with each sample scanned 32 times to ensure spectral accuracy, reproducibility, and an optimized signal-to-noise ratio. The resulting spectra were subsequently compared with standard reference spectra to further confirm the chemical composition and purity of the synthesised PIB-MNPs.

### Experimental Design

Supplementary Figure 1 shows a schematic representation of the experimental workflow for evaluating the toxicity and biological effects of polyisobutylene (PIB) polymer nanoparticles (PIB-MNPs) in *Drosophila melanogaster*. After synthesizing the PIB-MNPs, late-stage pupae from stock cultures were collected and maintained on standard cornmeal-agar medium at 25°C until eclosion. One-day-old adult flies of mixed sex were randomly assigned to agar-sucrose bottles (≥25 flies/bottle) for treatment exposure. Five groups of (n=3) replicates each were tested: PIB-MNPs at 1, 2, and 3 mg, an SDS-positive control, and a water control. Treatments were delivered as yeast paste (50 mg yeast + 50 µL dH₂O), with the respective PIB-MNP dose, SDS, or no added components, and were homogenized. Paste was placed at the corners of each bottle with agar-sucrose source (egg-laying medium). Media and treatments were refreshed every two days over a 21-day exposure period. Bottles were maintained at 25°C, 60–70% relative humidity, and under a 12 h light/dark cycle throughout.

### Survival Assay

Survival analysis was carried out alongside the same treatment bottles over the 21-day experimental period. A minimum of 40 flies were assessed across all five treatment groups. Mortality was recorded separately for males and females every two days, concurrent with the media and treatment replacement. Dead flies were counted and removed at each observation point. The significance of survival between the genotypes was assessed using the Log-rank (Mantel-Cox) test. Moreover, the Gehan–Breslow–Wilcoxon test was used to assess the differences in early death among the genotypes in the study. Statistical significance was set at *P* < 0.05. Early deaths, as accounted for in the Gehan–Breslow–Wilcoxon test, suggest that the flies are at risk due to genetic modifications affecting development, leading to premature aging (Gupta et al. 2025).

### Climbing Assay

Locomotor ability was assessed using a negative geotaxis (climbing) assay as described by Ranjan et al. (2024) with a slight modification. Briefly, male and female flies from each treatment group were separated without anesthesia and transferred into standard glass vials. Following a 1-minute acclimatization period, vials were sharply tapped three times to displace flies to the bottom, marking the start of the assay. The number of flies reaching or crossing the marked length within 10 seconds was recorded. Three trial replicates were performed per group, with a 1-minute recovery interval between trials, and all assays were conducted at a consistent time of day to control for circadian variation and under restricted overhead lighting to minimize phototactic interference.

### Fluorescence Imaging of the *Drosophila* Brain

Neurodegeneration was evaluated by quantifying brain vacuolization on days 5, 10, 15, and 21 (n > 3 brains/group). Flies were anaesthetized, and heads were excised and fixed in 4% paraformaldehyde (PFA) for 15–20 min. Brains were dissected in 1× PBS, post-fixed in fresh 4% PFA for 15–20 min, and washed in 1× PBS. Tissues were permeabilized sequentially in 0.3% and 1% PBTx (15 min each), then stained overnight at 4°C with <u>phalloidin and DAPI</u>, respectively. Following a 1% PBTx wash, brains were mounted and imaged at 10× magnification on an Olympus IX70 epifluorescence microscope. Vacuoles were manually counted across the entire brain and compared across treatment groups and each time point.

### Fertility and Fecundity Assay

Fertility and fecundity were assessed after 5 and 10 days of PIB-MNPs exposure using the method of El Kholy and Naggar (2023) with a slight modification. One-day-old virgin flies were then exposed to their respective treatments in agar-sucrose/yeast paste bottles, with media and treatment refreshed every 2 days.

For female fertility and fecundity, treated females were crossed individually with untreated 2-day-old *w¹¹¹⁸* males (1:1) for 12 h, then males were removed, and females were allowed to oviposit individually for 24 h. Eggs were counted to determine fecundity; plates were then incubated for a further 24 h, and hatched larvae were counted to determine fertility (percentage of eggs hatched). Ovaries were additionally dissected and examined at 10× magnification for the ratio of mature to immature egg chambers as an indicator of gonadal integrity.

For male fertility, treated males were crossed with untreated 2-day-old *w¹¹¹⁸* virgin females (1 male: 2 females) for 48 h. Males were then removed, and females were allowed to lay eggs and incubated for 7–10 days, with eclosed adult progeny counted as a measure of male fertility. Testes were dissected and examined at 10× magnification for the presence and proportion of late-stage cyst cells. All assays used a minimum of 6 flies per group per replicate.

### Smurf Assay

Gut barrier integrity was assessed on day 21 using the Smurf assay. From day 5 onward, blue food dye was incorporated into the yeast paste and agar-sucrose medium of all treatment groups and refreshed every 2 days until day 21 (n = 3 flies/group). Flies were anaesthetized and examined under a stereomicroscope (Olympus SZX12). Flies exhibiting dye leakage beyond the gut into the body cavity were classified as “Smurf” flies, indicative of increased intestinal permeability (Dambroise et al. 2016).

### Ellman’s Assay

Acetylcholinesterase (AChE) activity was measured on day 21 using the Ellman assay. Day 21 samples used only male heads due to reduced female survival. A minimum of 30 heads per group were homogenized on ice in 100 µL RIPA buffer containing protease inhibitors, DTT, and PMSF, then centrifuged (12,000 rpm, 20 min, 4°C), and the supernatant was stored at −80°C. Protein concentration was determined using the CB™ Protein Assay kit (Bio-Rad, Cat. # 786-012/786-012T/786-893) with a BSA standard curve and read at 595 nm (Infinite® M200 Pro, Tecan).

AChE activity was assayed using DTNB and acetylthiocholine iodide (ATCI) as chromogen and substrate. Samples were combined with 0.3 mM DTNB and incubated for 10 min before addition of 40 mM ATCI, with absorbance read at 412 nm at five sequential 1-minute intervals. Activity was calculated against a glutathione standard curve and normalized to protein concentration (mg/mL).

### RNA Extraction and RT-qPCR

Total RNA was extracted on day 21 using TRIzol (Sigma-Aldrich) reagent. Following standard chloroform-isopropanol extraction, RNA was resuspended in RNase-free water and stored at −80°C. RNA purity and integrity were confirmed by a NanoDrop ND-1000 spectrophotometer (Thermo Fisher Scientific) and 1% agarose gel electrophoresis for 28S/18S rRNA band integrity. cDNA was synthesized from 1 µg total RNA using a Bio-Rad reverse transcription kit (25°C, 46°C, 95°C, held at 4°C). Primers for *Stat92E* (target) and *rp49* (reference) were designed using Primer3 based on NCBI reference sequences and validated for specificity via Primer-BLAST. RT-qPCR reactions (10 µL total) contained 5 µL master mix, 0.5 µL each primer, 3 µL nuclease-free water, and 1 µL cDNA, run for 40 cycles with melt-curve analysis to confirm amplification specificity. PCR products were further verified by 1.5% agarose gel electrophoresis. Relative *Stat92E* expression was calculated using the 2^−ΔΔCt method, normalized to *rp49*, with the water control set as reference (fold change = 1). All assays were performed once across all treatment groups.

### Statistical Analysis

All quantitative data were analyzed using GraphPad Prism version 8.0 (GraphPad Software, San Diego, CA, USA). Data are presented as the mean ± standard error of the mean (SEM). Statistical comparisons among multiple experimental groups were performed using Student’s t-test. Relative gene expression was analyzed using the 2^−ΔΔCt method after normalization to the reference gene *rp49*, with the water control group serving as the calibrator. Differences were considered statistically significant at p < 0.05.

## Results

### Characterization of PIB-MNPs

#### a. Morphology characterization using Scanning Electron Microscopy (SEM)

The surface morphology and structural features of the synthesized PIB-MNPs were examined using a SEM (JEOL JSM-IT300) at two magnifications (×1,500 and ×7,000). The micrographs revealed that the synthesized PIB-MNPs exhibited irregular, flake-like morphology with heterogeneous size distribution across the particle population. At higher magnification (×7,000), particle size measurements indicated a range from the nanoscale to the microscale, with dimensions from approximately 382.62 nm to 7.174 µm, confirming the successful synthesis of particles within the micro/nanoplastic size range. At lower magnification (×1,500), the particles appeared as larger irregular aggregates with sizes ranging from approximately 2.586 µm to 18.59 µm, suggesting a tendency for particle aggregation **(Figures 1A and B).**

**Figure 1.**
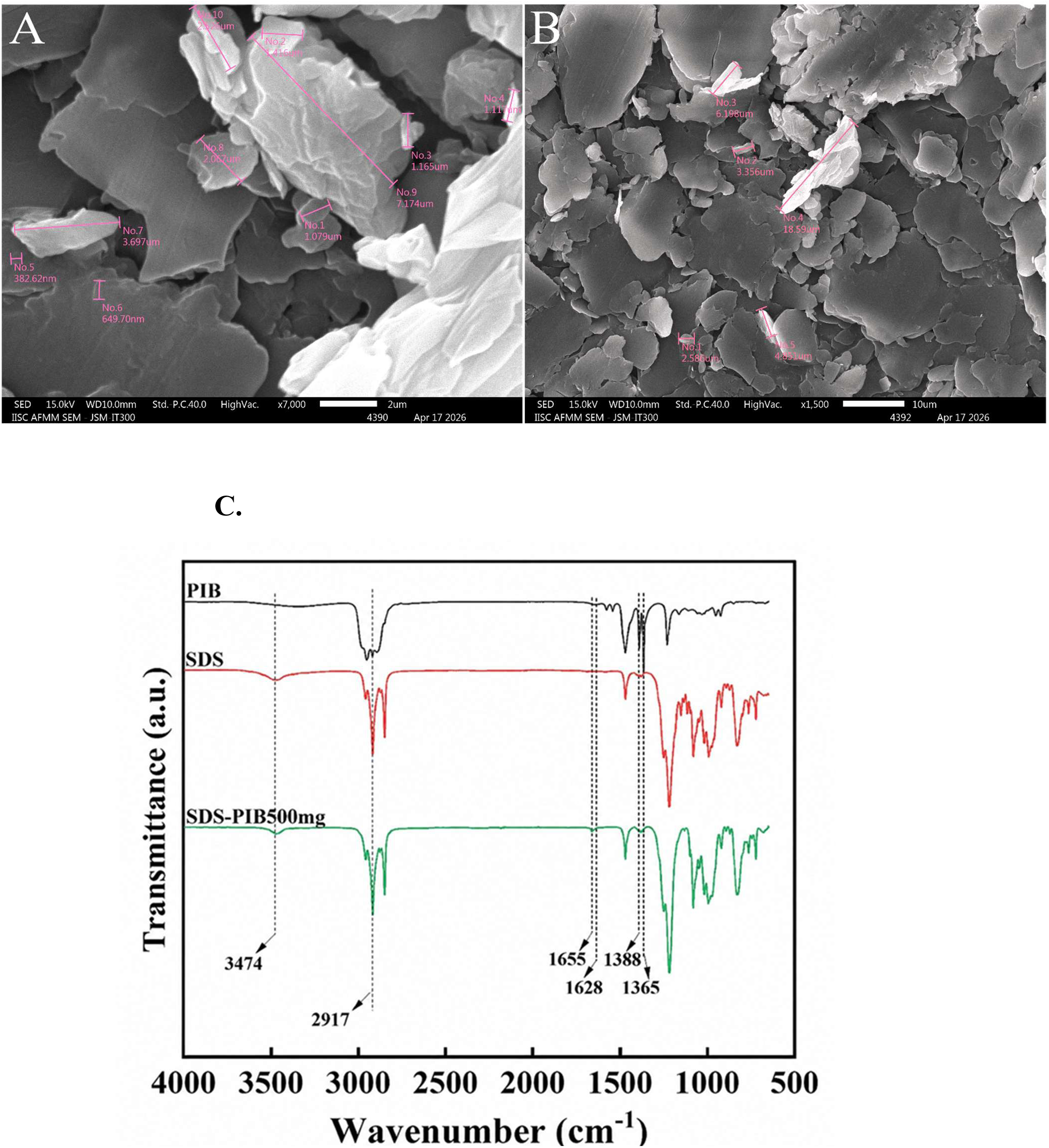
Scanning electron microscopy (SEM) characterization of PIB-MNPs. SEM micrographs of synthesized PIB-MNPs acquired using a JEOL JSM-IT300 (SED, 15.0 kV, WD 10.0 mm, HighVac) at **(A)** ×7,000 magnification (scale bar = 2 μm) and **(B)** ×1,500 magnification (scale bar = 10 μm). Representative particles are annotated (No. 1–10 in A; No. 1–5 in B) with corresponding dimensions, illustrating the irregular, flake-like morphology and heterogeneous size distribution of the particles across nano- to micro- scale ranges. Figure 1(C) shows FTIR spectra of pure PIB (top black), pure SDS (middle red), and PIB-MNPs (bottom green). Pure PIB exhibits characteristic C–H stretching at 2917 cm⁻¹ and a gem-dimethyl doublet at 1388/1365 cm⁻¹, diagnostic of the isobutylene repeat unit. Pure SDS shows an O–H stretch at 3474 cm⁻¹ along with S=O stretching bands from the sulfate head group. The PIB-MNP spectrum retains the diagnostic PIB bands (2917 and 1388/1365 cm⁻¹) and the SDS-associated O–H stretch (3474 cm⁻¹), while shifted absorptions at 1655 and 1628 cm⁻¹ indicate interfacial interaction between the PIB coating and the SDS-treated plastic particle surface.

#### b. Chemical properties characterization using Fourier Transform Infrared (FTIR) spectroscopy

The chemical identity and functional group composition of the synthesized PIB-MNPs were confirmed by FTIR spectroscopy in ATR mode using a PerkinElmer Spectrum One system. The FTIR spectra of pristine PIB, SDS, and the synthesized PIB-MNPs were compared to identify characteristic peaks and confirm the chemical nature of the synthesized particles. The spectrum of the synthesized PIB-MNPs displayed a broad absorption band at 3474 cm⁻¹ and a strong absorption peak at 2917 cm⁻¹, both of which correspond to characteristic peaks of SDS, confirming the presence of residual SDS in the synthesized PIB-MNPs. Absorption bands at 1655 cm⁻¹ and 1628 cm⁻¹ were characteristic peaks of the PIB backbone, while peaks at 1388 cm⁻¹ and 1365 cm⁻¹ corresponded to characteristic gem-dimethyl deformation bands, which are a hallmark of the polyisobutylene structure (Figure 1C). The overall spectral profile of the synthesized PIB-MNPs showed concordance with both the reference PIB and SDS spectra, with the presence of characteristic peaks from both components collectively confirming the successful synthesis of PIB-MNPs with SDS as the surfactant stabilizer, as mentioned in previous studies (Anifowoshe et al. 2024).

### Chronic PIB-MNPs Exposure Compromises Survival of D. melanogaster with Greater Susceptibility Observed in Females

The effect of PIB-MNPs exposure on the survival of *D. melanogaster* was assessed over a 21-day period across all five treatment groups and recorded separately for male and female flies (Figure 12). Overall, a declining trend in percent survival was observed in PIB-MNPs treated flies relative to control groups in both sexes, with females exhibiting a markedly accelerated rate of mortality compared to males throughout the experimental period.

In female flies (Figure 2A), survival declined rapidly across all PIB-MNPs treatment groups from the early stages of exposure. The 1 mg, 2 mg, and 3 mg PIB-MNPs groups showed a steep and progressive decline in survival, with near-complete mortality observed by day 21 in all three treatment groups. The SDS control group also exhibited a notable decline in survival relative to the water control, suggesting a degree of SDS-associated toxicity at the concentration used. The water-control females maintained comparatively higher survival throughout the 21-day period, recording approximately 22% survival at day 21.

**Figure 2:**
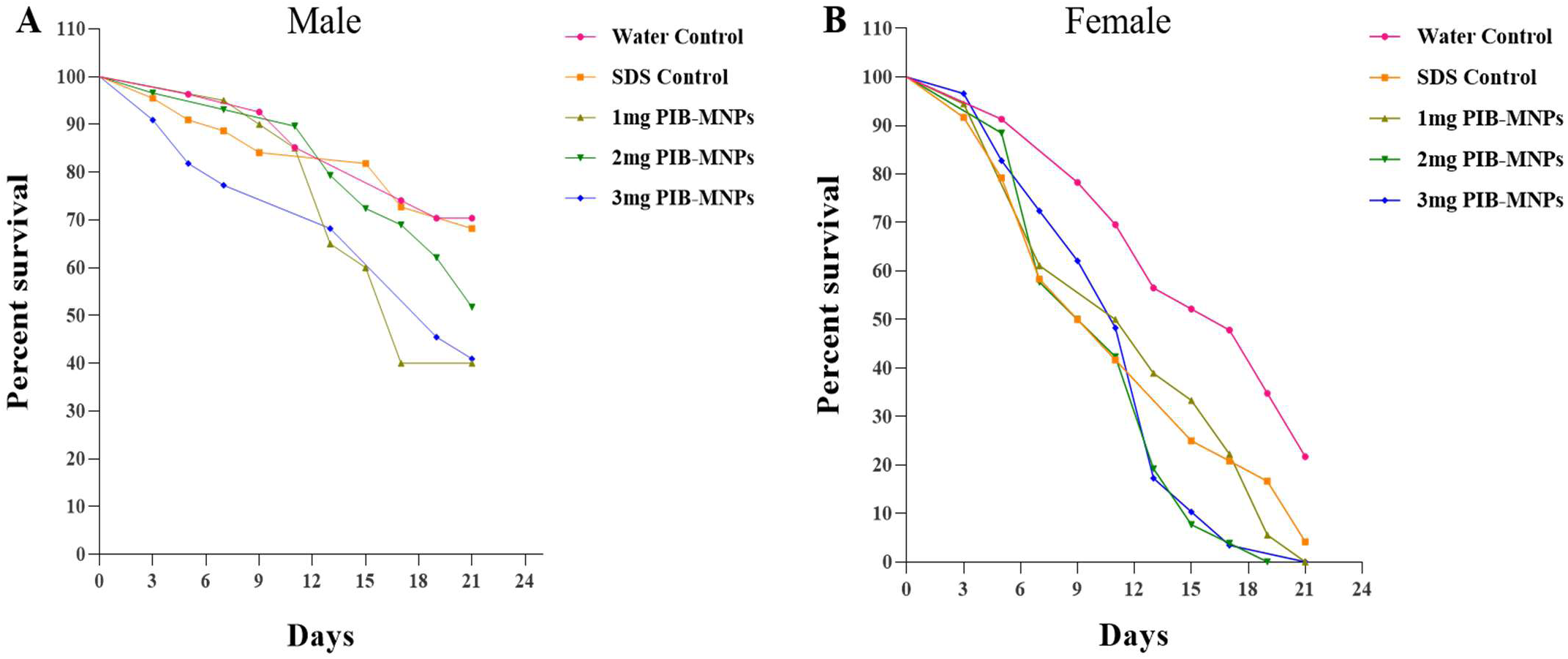
Kaplan-Meier survival curves showing percent survival over time (days) in male (A) and female (B) *D. melanogaster* exposed to PIB-MNPs over 21 days. Percent survival was recorded every two days across all five treatment groups (water control, SDS control, 1 mg, 2 mg, and 3 mg PIB-MNPs) throughout the 21-day experimental period. Female flies exhibited a rapid and progressive decline, while male flies showed a more gradual decline in survival.

In male flies (Figure 2B), survival declined more gradually across all treatment groups compared to females. The water-control males maintained the highest survival throughout the experimental period, recording approximately 69% survival at day 21. The SDS control group showed a moderate decline in survival relative to the water control. Among the PIB-MNPs treatment groups, a declining trend in survival was observed across all concentrations, with the 1 mg and 2 mg PIB-MNPs groups recording approximately 40% survival at day 21, while the 3 mg PIB-MNPs group recorded approximately 51% survival at the same time point. The comparatively higher survival observed in male flies across all treatment groups relative to females suggests a sex-dependent and concentration-independent difference in sensitivity to PIB-MNPs exposure.

### PIB-MNPs Exposure Impairs Locomotor Activity in D. melanogaster in a Dose-Independent and Time-Dependent Manner

Locomotor function was assessed using a negative geotaxis (climbing) assay, with performance expressed as the percentage of flies crossing an 8 cm mark within 10 s. Female flies were assessed on days 5 and 10, and male flies on days 5, 10, 15, and 21 of PIB-MNP exposure (Figure 3).

**Figure 3.**
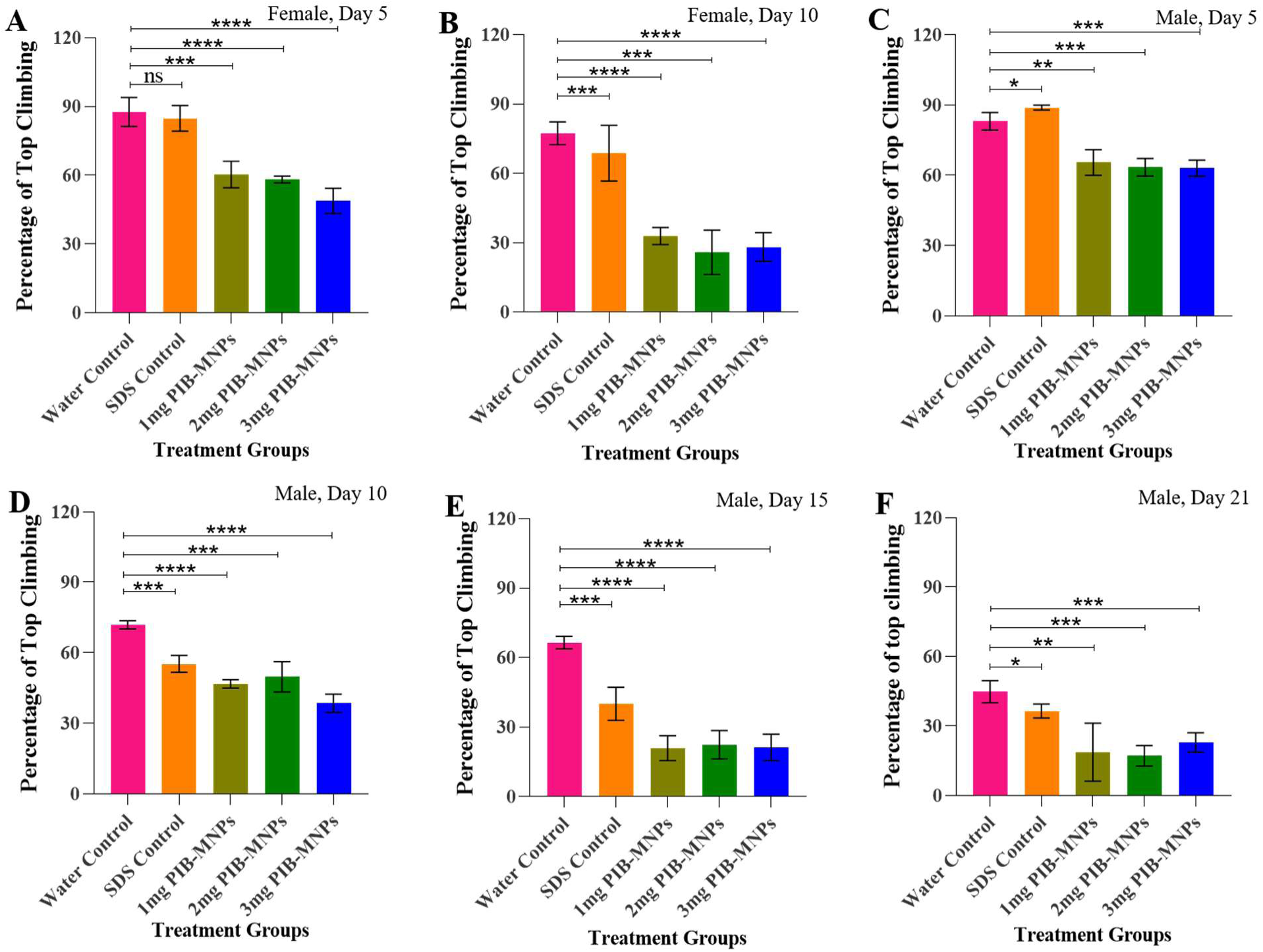
PIB-MNP exposure impairs climbing performance in *D. melanogaster* in a dose-independent, time-progressive manner. Locomotor activity was assessed by the negative geotaxis (climbing) assay and expressed as the percentage of flies crossing an 8 cm mark within 10 s. Female flies were assessed at (A) day 5 and (B) day 10, and male flies at (C) day 5, (D) day 10, (E) day 15, and (F) day 21 of exposure to water control, SDS control, or 1, 2, or 3 mg PIB-MNPs. Data are presented as mean ± SEM (n > 20 flies/replicate, 3 replicates/group). Statistical significance was determined by Student’s t-test relative to the water control; ns = not significant, *p < 0.05, **p < 0.01, ***p < 0.001, ****p < 0.0001.

In females (Figures 3A, B), climbing performance declined markedly in all PIB-MNP treatment groups relative to both controls at both time points, with the magnitude of decline not scaling with dose. On day 5 (Figure 3A), water and SDS controls achieved comparable climbing rates of ∼88% and ∼85%, respectively, while the 1, 2, and 3 mg PIB-MNP groups fell to ∼60%, 58%, and 49%. By day 10 (Figure 3B), overall performance had declined across all groups—water and SDS controls reached ∼77% and 68%—with PIB-MNP-treated flies more severely impaired at ∼32%, 26%, and 28% for the 1, 2, and 3 mg groups.

Males (Figures 3C–F) showed a similar pattern of impairment that became progressively more pronounced with prolonged exposure. On day 5 (Figure 3C), water and SDS controls climbed at ∼83% and 89%, while the 1, 2, and 3 mg PIB-MNP groups declined to ∼65%, 63%, and 63%. On day 10 (Figure 3D), controls dropped to ∼72% (water) and 55% (SDS), with treated groups further reduced to ∼47%, 50%, and 39%. By day 15 (Figure 3E), the water control declined to ∼66% and the SDS control to ∼40%, while the PIB-MNP groups fell sharply to ∼21%, 23%, and 22%. At day 21 (Figure 3F), the water control had dropped to ∼44% and the SDS control to ∼36%, with treated groups further reduced to ∼18%, 17%, and 23%.

Taken together, these results indicate that chronic PIB-MNP exposure progressively impairs locomotor function in both sexes, independent of dose, with male flies showing a consistent, cumulative decline across all four assessment points over the 21-day exposure period. Notably, climbing performance in the control groups also declined over time in males, consistent with age-related locomotor decline reported in *D. melanogaster*, but PIB-MNP-treated flies remained significantly more impaired than age-matched controls at every time point.

### Chronic PIB-MNPs Exposure Induces Progressive Brain Vacuolization in D. melanogaster

Brain vacuolization was examined by quantifying vacuoles in phalloidin/DAPI-stained whole-mount brain preparations at days 5, 10, 15, and 21 of PIB-MNPs exposure (Figure 4). Vacuoles appeared as distinct unstained void regions within the brain parenchyma, marked by white arrows in the representative images (Figure 4, A–D). Across all four time points, the water and SDS control groups showed minimal vacuole formation, remaining below 2–3 vacuoles per brain, while the PIB-MNPs-treated groups showed a consistent increase in vacuole number over the course of exposure, irrespective of concentration.

**Figure 4.**
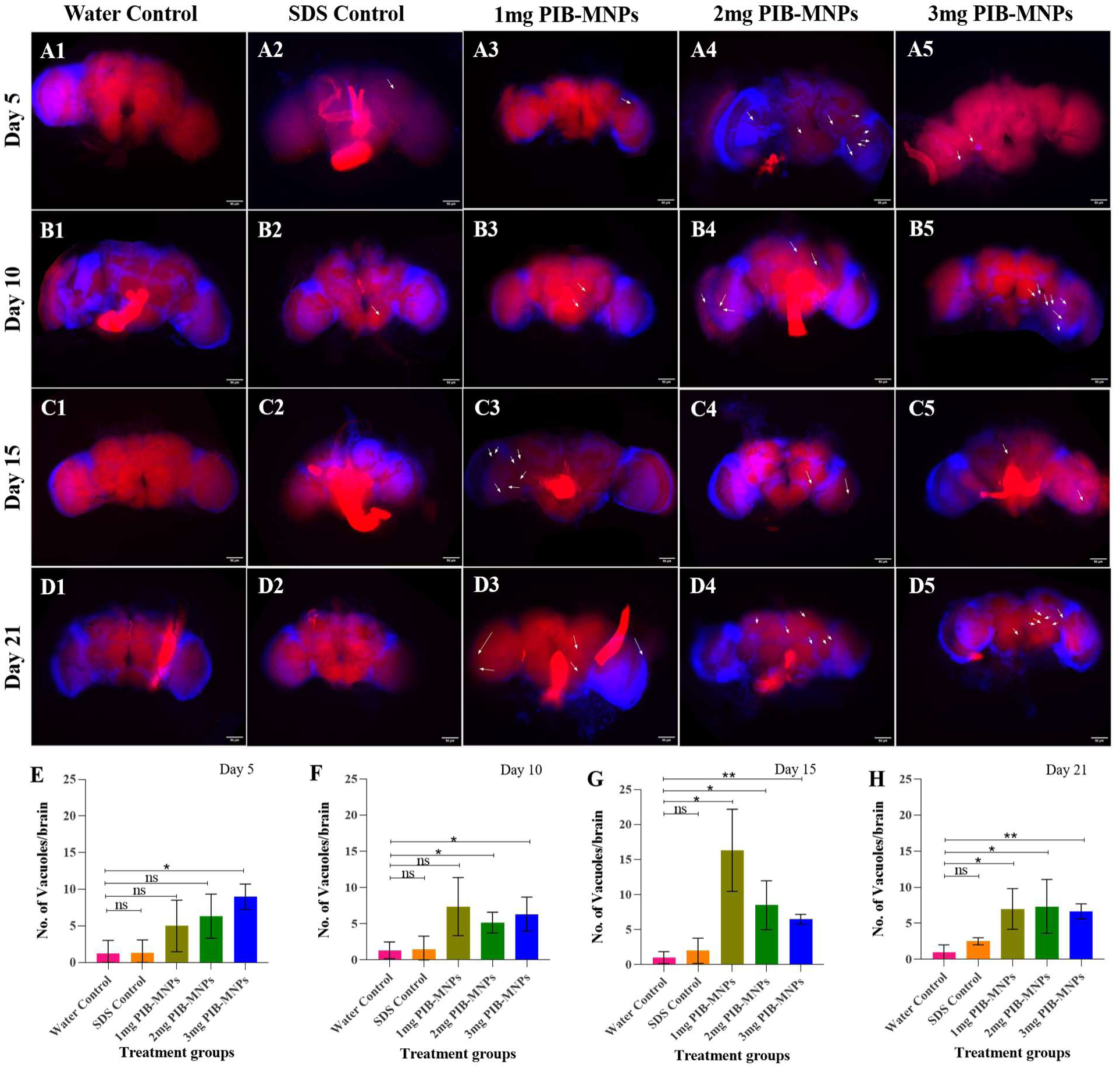
Representative fluorescence micrographs of *Drosophila melanogaster* brains. (phalloidin/DAPI-stained, red/blue merge) across treatment groups and exposure durations, with quantification of vacuole formation. (A–D) Whole-mount brain images from flies exposed to Water Control, SDS Control, 1 mg, 2 mg, and 3 mg PIB-MNPs (columns, left to right) at Day 5 (A1–A5), Day 10 (B1–B5), Day 15 (C1–C5), and Day 21 (D1–D5) post-exposure. White arrows indicate vacuolar lesions within the brain. Scale bar = 50 μm. (E–H) Quantification of the number of vacuoles per brain at Day 5 (E), Day 10 (F), Day 15 (G), and Day 21 (H) across the five treatment groups. Data are presented as mean ± SD (n > 5 brains/group, N=3). Statistical significance was determined by Student’s *t*-test, with each treatment group compared to the water control group (ns = not significant, *p < 0.05, **p < 0.01).

At day 5, vacuole numbers remained low across all groups, with the 1, 2, and 3 mg PIB-MNPs groups recording approximately 5, 6, and 9 vacuoles per brain, respectively. Only the 3 mg group differed significantly from the water control (p < 0.05), while the 1 mg and 2 mg groups showed an upward trend that did not reach statistical significance (Figure 4, E). By day 10, the 1 mg and 3 mg PIB-MNPs groups showed significant increases over the water control (p < 0.05), recording approximately 7 and 6 vacuoles per brain, respectively, while the 2 mg group (∼5 vacuoles/brain) did not differ significantly from the control (Figure 4, F).

The most pronounced effect was observed at day 15, where the 1 mg PIB-MNPs group recorded the highest vacuole count observed across the entire study, at approximately 16 vacuoles per brain (p < 0.05), exceeding both the 2 mg (∼8 vacuoles/brain, p < 0.01) and 3 mg (∼6 vacuoles/brain, p < 0.01) groups (Figure 4, G). This pattern indicates that vacuole accumulation did not scale consistently with PIB-MNPs concentration and that the lowest dose tested produced the greatest effect. By day 21, vacuole numbers across all three PIB-MNPs groups converged to a comparable range of approximately 7 vacuoles per brain, remaining significantly elevated relative to the water control (1 mg: p < 0.05; 2 mg and 3 mg: p < 0.01) (Figure 4, H). These results indicate that chronic PIB-MNPs exposure induces a time-dependent, but not strictly dose-dependent, increase in brain vacuolization, with the highest vacuole numbers observed at day 15 across all treated groups.

### PIB-MNPs Exposure Is Associated with Increased Intestinal Barrier Permeability

Intestinal barrier integrity was evaluated using the Smurf assay at day 21 of exposure. Flies were scored as “Smurf” positive based on the visible spread of blue dye beyond the gut into the body cavity, assessed under a stereomicroscope (Supplementary Figure 4). Water control flies (Supplementary Figure 4A) showed dye largely confined to the gut, with minimal leakage into the body cavity. Flies from the 1, 2, and 3 mg PIB-MNPs groups (Supplementary Figure 4C–E) showed a broader distribution of dye within the body cavity, consistent with increased gut permeability relative to the water control. The SDS control (Supplementary Figure 4B) also displayed some visible dye spread beyond the gut. As all flies were assessed at 21 days of age, age-associated increases in gut permeability may have contributed to the observed phenotype independently of PIB-MNPs exposure. This assay was performed qualitatively, based on visual assessment of representative images (N = 2, n = 3 flies/group); no quantitative scoring of Smurf-positive frequency was carried out.

### PIB-MNPs Exposure Affects Fertility and Fecundity in D. melanogaster with Associated Gonadal Disruption

#### a. PIB-MNPs Exposure Reduces Egg Laying and Hatching Rate in Female D. melanogaster

Female fecundity and fertility were assessed at days 5 and 10 of PIB-MNPs exposure by recording the number of eggs laid per female and the corresponding hatching rate (Figure 5). At day 5, the water control and SDS control recorded approximately 25 and 18 eggs laid per female, respectively, while the 1 mg, 2 mg, and 3 mg PIB-MNPs groups recorded approximately 7, 8, and 6 eggs per female, respectively, representing a clear reduction in egg laying across all treatment groups compared to the water control (1 mg and 3 mg: p < 0.001; 2 mg: p < 0.01) (Figure 5, D). The hatching rate at day 5 followed a similar declining pattern, with the water control and SDS control recording approximately 90% and 80% hatching, respectively, compared to approximately 55%, 20%, and 58% in the 1 mg, 2 mg, and 3 mg PIB-MNPs groups, respectively. Of these, only the 2 mg and 3 mg groups differed significantly from the water control (p < 0.01 and p < 0.05, respectively), while the 1 mg group did not reach statistical significance (Figure 5, E).

**Figure 5.**
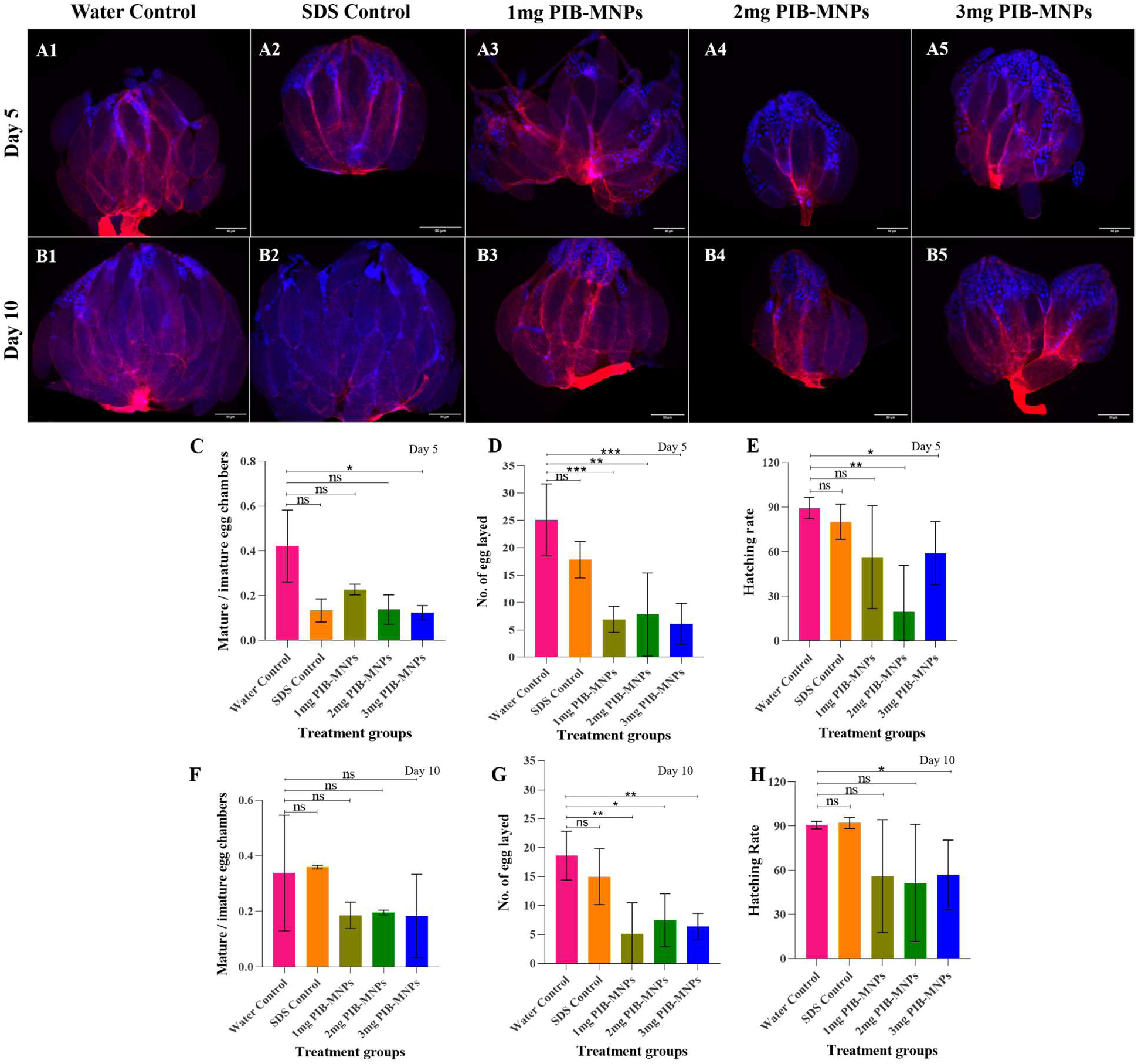
Effect of PIB-MNPs exposure on female fertility, fecundity, and ovarian egg chamber maturation in *D. melanogaster*. (A, B) Representative epifluorescence micrographs of dissected ovaries stained with phalloidin (red) and DAPI (blue) from water control, SDS Control, 1 mg, 2 mg, and 3 mg PIB-MNPs treatment groups (columns, left to right) at day 5 (A1–A5) and day 10 (B1–B5) of exposure. Scale bar = 50 μm. (C, F) Quantification of the mature-to-immature egg chamber ratio at day 5 (C) and day 10 (F) across treatment groups. (D, G) Number of eggs laid per female at day 5 (D) and day 10 (G) across treatment groups. (E, H) Hatching rate (%) at day 5 (E) and day 10 (H) across treatment groups. Data are presented as mean ± SEM (n ≥ 5 flies/group, N = 2). Statistical significance was determined using Student’s *t*-test, with each treatment group compared to the water control group (ns = not significant, *p < 0.05, **p < 0.01, ***p < 0.001).

At day 10, egg-laying values in the water control and SDS control were approximately 18 and 15 eggs per female, respectively, while the 1 mg, 2 mg, and 3 mg PIB-MNPs groups recorded approximately 5, 7, and 6 eggs per female, respectively, remaining significantly reduced relative to the water control (1 mg and 3 mg: p < 0.01; 2 mg: p < 0.05) (Figure 5, G). The hatching rate on day 10 was approximately 90% and 92% in the water and SDS controls, respectively, compared to approximately 55%, 50%, and 57% in the 1 mg, 2 mg, and 3 mg PIB-MNPs groups, respectively. Only the 3 mg group showed a statistically significant reduction relative to the water control (p < 0.05), while the 1 mg and 2 mg groups did not differ significantly, likely reflecting the variability observed within these groups (Figure 5, H). Overall, a consistent decline in both egg-laying and hatching rates was observed across all PIB-MNPs treatment groups at both time points, indicating an adverse effect of PIB-MNPs exposure on female fecundity and fertility.

#### b. PIB-MNPs Exposure Disrupts Ovarian Egg Chamber Maturation in Female D. melanogaster

The effect of PIB-MNPs exposure on female gonadal integrity was examined by determining the ratio of mature to immature egg chambers in dissected ovaries at days 5 and 10 of exposure (Figure 5 A and B). At day 5, water control ovaries showed a mature-to-immature egg chamber ratio of approximately 0.42, whereas the SDS control, 1 mg, 2 mg, and 3 mg PIB-MNPs groups recorded ratios of approximately 0.13, 0.23, 0.14, and 0.12, respectively. Only the 3 mg group differed significantly from the water control (p < 0.05), while the remaining groups showed a downward trend that did not reach statistical significance (Figure 5, C).

At day 10, the water control and SDS control maintained comparable ratios of approximately 0.34 and 0.36, respectively, while the 1 mg, 2 mg, and 3 mg PIB-MNPs groups recorded ratios of approximately 0.19, 0.20, and 0.18, respectively. None of the treatment groups differed significantly from the water control at this time point, although all three PIB-MNPs groups continued to show a lower mean ratio, consistent with the trend observed at day 5 (Figure 5, F). Representative micrographs of dissected ovaries showed a visibly reduced proportion of mature egg chambers and a correspondingly higher proportion of immature chambers in the PIB-MNPs-treated groups relative to controls, in line with the quantified reduction in the mature-to-immature ratio.

#### a. PIB-MNPs Exposure Reduces the Number of Flies Eclosed

The effect of PIB-MNPs exposure on offspring emergence was assessed by quantifying the number of flies eclosed at days 5 and 10 of exposure (**Figure 6, C and E**). At day 5, the water control recorded approximately 85 flies eclosed, while the SDS control and 1 mg PIB-MNPs group recorded approximately 67 and 70 flies eclosed, respectively, neither of which differed significantly from the water control. In contrast, the 2 mg and 3 mg PIB-MNPs groups showed a significant reduction in the number of flies eclosed, recording approximately 61 and 54 flies, respectively (p < 0.01 for both), indicating a decline in offspring emergence.

**Figure 6.**
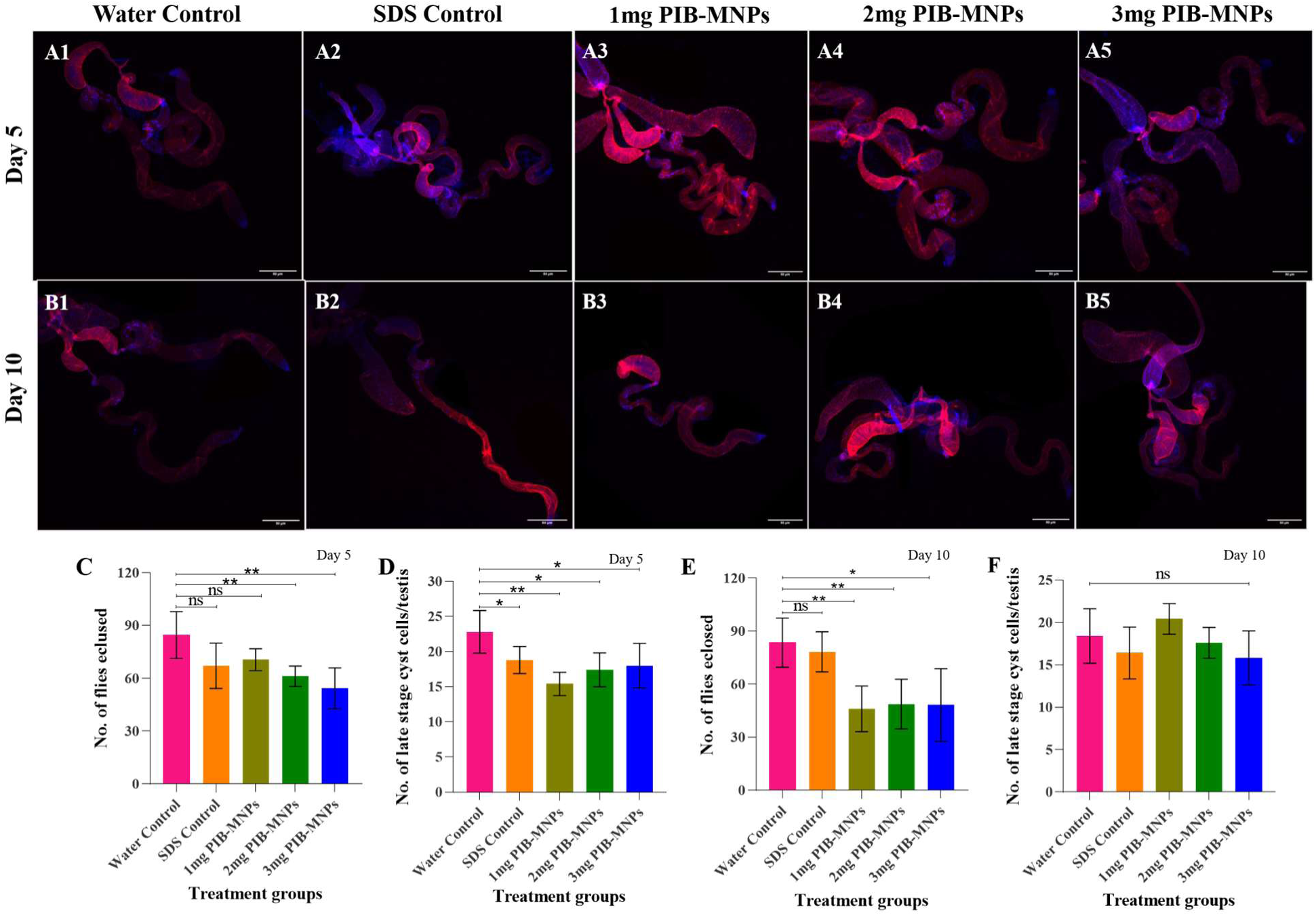
Effect of PIB-MNPs exposure on offspring emergence and altered nuclear density in the late-stage testis in male *D. melanogaster* testes. (A, **B)** Representative epifluorescence micrographs of dissected testes stained with phalloidin (red) and DAPI (blue) from Water Control, SDS Control, 1 mg, 2 mg, and 3 mg PIB-MNPs treatment groups (columns, left to right) at day 5 **(A1–A5)** and day 10 **(B1–B5)** of exposure. Scale bar = 50 μm. **(C, E)** Number of flies eclosed at day 5 **(C)** and day 10 **(E)** across treatment groups. **(D, F)** Number of late-stage cyst cells per testis at day 5 **(D)** and day 10 **(F)** across treatment groups. Data are presented as mean ± SEM (n ≥ 5 flies/group, N = 2). Statistical significance was determined using Student’s *t*-test, with each treatment group compared to the water control group (ns = not significant, *p < 0.05, **p < 0.01).

At day 10, the water control and SDS control recorded approximately 83 and 78 flies eclosed, respectively, with no significant difference between the two. All three PIB-MNPs treatment groups, however, showed a marked reduction relative to the water control, with the 1 mg, 2 mg, and 3 mg groups recording approximately 46, 48, and 48 flies eclosed, respectively (1 mg: p < 0.01; 2 mg and 3 mg: p < 0.05). Taken together, these results indicate that PIB-MNPs exposure reduces the number of flies eclosed, with the effect becoming more pronounced and consistent across all treatment groups by day 10.

#### b. PIB-MNPs Exposure Shows No Notable Effect on altered nuclear density in the late-stage testis in Male D. melanogaster Testes

The effect of PIB-MNPs exposure on male gonadal integrity was assessed by quantifying the number of late-stage cyst cells per testis in dissected testes at days 5 and 10 of exposure (**Figure 6, D and F**). At day 5, the water control recorded approximately 23 late-stage cyst cells per testis, while the SDS control and the 1 mg, 2 mg, and 3 mg PIB-MNPs groups recorded approximately 19, 16, 17, and 19 late-stage cyst cells per testis, respectively, each differing significantly from the water control (SDS control and 3 mg: p < 0.05; 1 mg and 2 mg: p < 0.01).

At day 10, the water control recorded approximately 19 late-stage cyst cells per testis, while the SDS control and the 1 mg, 2 mg, and 3 mg PIB-MNPs groups recorded approximately 16, 20, 17, and 16 late-stage cyst cells per testis, respectively, none of which differed significantly from the water control. Overall, although a significant reduction in late-stage cyst cell number was observed across treatment groups at day 5, this effect was not sustained at day 10, and no consistent declining trend was evident across the two time points. These findings suggest that PIB-MNPs exposure at the concentrations tested did not substantially or persistently impair spermatogenesis, as assessed by altered nuclear density in the late-stage testis, despite its effect on the number of flies eclosed.

### PIB-MNPs Exposure Reduces Acetylcholinesterase Activity

Acetylcholinesterase (AChE) activity was assessed using Ellman’s assay by monitoring the change in absorbance at 412 nm over a 15-minute period (Figure 7A). The water control recorded absorbance values ranging from approximately 0.88 at minute 0 to 0.95 at minute 15, reflecting a steady increase indicative of active enzymatic hydrolysis of the substrate. The SDS control recorded absorbance values ranging from approximately 1.04 to 1.06 across the same time period, showing only a marginal increase relative to the water control. The 1 mg PIB-MNPs group recorded absorbance values ranging from approximately 0.68 to 0.71, showing a similarly limited increase over time. The 2 mg PIB-MNPs group recorded absorbance values ranging from approximately 0.55 to 0.55, remaining essentially unchanged throughout the assay, while the 3 mg PIB-MNPs group recorded absorbance values ranging from approximately 1.18 to 1.19, also showing no appreciable change across the 15-minute period. Overall, these results indicate that PIB-MNPs exposure reduces AChE activity, with the 2 mg and 3 mg groups showing the most pronounced loss of enzymatic activity.

**Figure 7.**
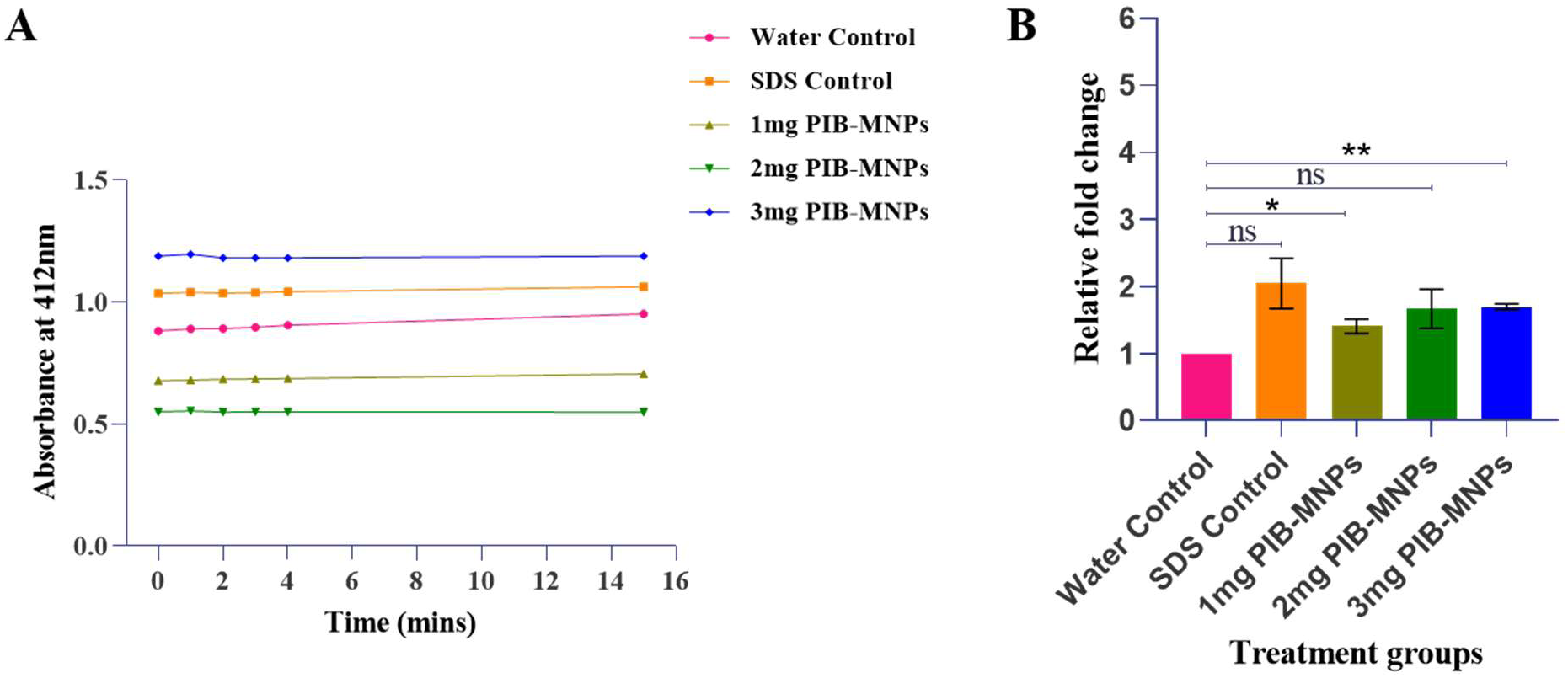
Effect of PIB-MNPs exposure on acetylcholinesterase (AChE) activity and *Stat92E* gene expression in *D. melanogaster*. **(A)** AChE activity assessed by Ellman’s assay, expressed as absorbance at 412 nm measured over a 15-minute period, across Water Control, SDS Control, 1 mg, 2 mg, and 3 mg PIB-MNPs treatment groups. **(B)** Relative fold change in *Stat92E* gene expression, determined by RT-qPCR and normalized to the Water Control group, across all treatment groups. Data are presented as mean ± SEM. Statistical significance was determined using Student’s *t*-test, with each treatment group compared to the water control group (ns = not significant, *p < 0.05; **p < 0.01; N = 2).

### PIB-MNPs Exposure Upregulates Stat92E Gene Expression

To determine whether AChE activity changes were accompanied by broader stress- or immune-related transcriptional changes, relative *Stat92E* gene expression, a marker of JAK/STAT pathway activation associated with immune and stress responses in *Drosophila* (Myllymäki, H. et al., 2014) was quantified by RT-qPCR across all treatment groups (Figure 7, B). Relative to the water control (set at a fold change of 1), the SDS control, 1 mg, 2 mg, and 3 mg PIB-MNPs groups showed fold changes of approximately 2.05, 1.4, 1.7, and 1.7, respectively. Of these, the 1 mg and 3 mg PIB-MNPs groups showed a statistically significant increase in *Stat92E* expression relative to the water control (p < 0.05 and p < 0.01, respectively), while the SDS control and 2 mg PIB-MNPs group did not differ significantly. Taken together, these results indicate that PIB-MNPs exposure leads to an upregulation of *Stat92E* gene expression, consistent with activation of the JAK/STAT signaling pathway.

## Discussion

The physicochemical characterization of synthesized PIB-MNPs confirmed retention of polyisobutylene’s chemical features with residual surfactant from emulsification. FTIR revealed gem-dimethyl deformation bands of the PIB backbone alongside sodium dodecyl sulfate (SDS) peaks, indicating surfactant traces remained. SDS acts as a surfactant during microplastic synthesis, stabilizing emulsions, reducing interfacial tension, and preventing particle coalescence, thereby enabling controlled formation of uniform polymeric microplastic particles. Thus, our SEM results showed irregular morphology with aggregated structures below 5 µm. Similar findings were reported by Anifowoshe et al. (2024) in zebrafish studies employing SDS-assisted emulsification, where the synthesis and characterization of micro-sized polyisobutylene and its toxicological effects on zebrafish *(Danio rerio*) development and homeostasis were investigated. Beyond vertebrate models such as zebrafish, *Drosophila* has also demonstrated strong utility in revealing the hazardous impacts of nanoscale materials across multiple studies (Alaraby et al., 2016). Thus, to investigate the potential adverse effects of polyisobutylene micro- and nanoplastics (PIB-MNPs), we employed *Drosophila melanogaster* (*w^1118^*) as a robust and genetically tractable experimental model. This model enabled the evaluation of PIB-MNP-induced toxicological outcomes across multiple biological endpoints, thereby providing mechanistic insights into the impacts of these emerging plastic pollutants.

Chronic dietary exposure to PIB-MNPs in *Drosophila* reduced survival, with females showing greater susceptibility than males across 21 days, approaching complete mortality by day 21. Sex-specific vulnerability to contaminants has been linked to metabolic activity, reproductive investment, and antioxidant capacity (Jin et al., 2021; Sharma & Chatterjee, 2017). Mortality did not increase proportionally with concentration; the 3 mg group was not consistently more toxic than lower doses, reflecting non-monotonic responses reported in microplastic toxicology (Lenz et al., 2016; Hwang et al., 2020). Aggregation reduced bioavailability, explaining comparable effects at lower doses. SDS controls caused intermediate mortality, but PIB-MNPs remained the principal toxicant.

Next, we conducted a behavioral climbing assay. Declining climbing ability emerged as a clear marker of chronic neurological impairment, with males exhibiting a gradual reduction in locomotor performance from day 5 to day 21 across all concentrations, indicating negative geotaxis. Negative geotaxis in *D. melanogaster* refers to the innate tendency of flies to orient and move against the force of gravity, typically by climbing upward when placed at the bottom of a vertical surface (Liao et al. 2012). This instinctive behavior is considered a robust and evolutionarily conserved response, enabling the flies to avoid unfavorable low-lying environments and seek elevated positions. It is widely employed in climbing assays as a reliable measure of locomotor performance, neuromuscular coordination, and overall vitality. Because the assay is simple, reproducible, and sensitive to subtle physiological changes, it has become a standard tool in *Drosophila* research for assessing age-related decline, genetic mutations affecting motor function, and the impact of environmental or toxicological stressors on locomotor ability (Ranjan et al., 2024). Thus, the pattern observed in this present study suggests interference with neuronal function through cumulative physiological damage rather than acute toxicity. Comparable reductions in locomotor activity have been documented in zebrafish and *Drosophila* exposed to polystyrene and polyethylene terephthalate microplastics, attributed to oxidative stress, mitochondrial dysfunction, and disrupted neurotransmission (Chen et al., 2017; Liu et al., 2022; Kauts et al., 2023). Neurobehavioral impairment thus appears to be a conserved response to chronic microplastic exposure. This result aligns with studies from Ranjan et al. (2024) and Raj et al. (2024). Similarly, our findings are consistent with those reported by Anifowoshe et al. (2024). In their behavioral assessments, including the novel tank and social interaction tests, distinct alterations in locomotor activity and exploratory behavior were observed between the control group and flies exposed to varying concentrations of PIB-MPs. The reduction in locomotor activity noted in our study parallels evidence from murine models, where polystyrene microplastic–induced overproduction of reactive oxygen species (ROS) disrupted skeletal muscle regeneration by altering satellite cell fate (Shengchen et al., 2021). Therefore, the results indicate that PIB-MNPs exposure may have accumulated within the flies’ tissues, disrupting neuronal signaling and muscular integrity. This progressive physiological stress reduces mobility and strength, thereby impairing their instinctive climbing (negative geotaxis) response.

Microplastics (MPs) have been shown to disrupt the composition and diversity of gut microbial communities, leading to gut microbiota dysbiosis (Qiao et al., 2019). Given the critical role of the gut microbiome in regulating neurodevelopment, neurotransmission, immune signaling, and brain function through the gut-brain axis, such microbial perturbations may have far-reaching neurological consequences. Indeed, growing evidence implicates gut microbiome dysregulation in the pathogenesis of various neurological and neuropsychiatric disorders (Cryan et al., 2020).

Our next focus was on neurodegeneration. In these results, structural observations from whole-brain imaging of flies further reinforced the interpretation we observed in the climbing experiments. Brain vacuolization increased markedly in PIB-MNPs–treated flies, reaching its highest levels after 15 days of exposure. In *Drosophila*, vacuole formation is widely recognized as a histological hallmark of neurodegeneration, reflecting progressive neuronal loss and tissue deterioration (Behnke et al., 2021). The temporal pattern closely paralleled the decline in locomotor performance, suggesting that behavioural impairment was accompanied by structural deterioration of nervous tissue. The modest reduction in vacuole counts at the final sampling point is unlikely to indicate recovery. Instead, it likely reflects survivorship bias, as individuals with the most extensive neurodegeneration may not have survived to day 21. Consequently, the remaining flies represented a relatively less affected population despite prolonged exposure. Similar considerations have been highlighted in chronic toxicological studies, where substantial mortality alters cohort characteristics and complicates interpretation of terminal endpoints.

Because brain and gut functions are bidirectionally linked, we next examined gut integrity to correlate the neurological effects with intestinal physiology in *Drosophila* using Smurf assay. PIB-MNPs exposure produced a positive Smurf phenotype after 21 days, marked by dye leakage into the body cavity, indicating barrier dysfunction (Rera et al., 2012). This phenotype suggests compromised intestinal homeostasis, consistent with chronic inflammation, metabolic imbalance, and reduced lifespan. Similar effects have been reported in vertebrates, where polystyrene microplastics disrupted barrier integrity, altered microbiota, and disturbed hepatic metabolism (Lu et al., 2018; Hwang et al., 2020). Although age-related decline and qualitative assay limitations warrant caution, the findings support that prolonged PIB-MNPs exposure impairs epithelial integrity and intestinal physiology, aligning with the reports from Anifowoshe et al. (2024). Our study is similar to Alaraby et al. (2023), who reported that the internalization ability of PETNPL was associated with significant alterations in gene expression linked to general stress responses, oxidative damage, and intestinal barrier disruption.

In the reproductive toxicity experiments, female flies showed the greatest sensitivity to chronic PIB-MNPs exposure, with fecundity and egg hatching success declining markedly, especially after prolonged treatment. Fluorescence staining revealed fewer mature egg chambers, indicating disrupted ovarian development and oocyte maturation. As reproduction is energetically demanding in *Drosophila*, disturbances in nutrient allocation, endocrine regulation, or cellular homeostasis rapidly reduce egg output. The pronounced impairment aligns with female susceptibility evident in survival analysis. Comparable reproductive defects have been reported with polystyrene microplastics in mammals (Sharma & Chatterjee, 2017; Jin et al., 2021) and *Drosophila* (Liu et al., 2022), suggesting reproductive dysfunction is a conserved response to chronic MPs exposure.

Interestingly, male reproductive imaging showed comparatively limited structural alterations. Quantification of nuclear density in the late-stage testis revealed no obvious differences between treatment groups at either sampling point, suggesting that terminal spermatogenesis remained largely preserved under the exposure conditions employed in this study. This finding is consistent with previous reports demonstrating that PET microplastics (PET-MPs) accumulate in testicular tissue, leading to impaired sperm quality and reduced reproductive hormone levels in mice (Zhang et al., 2025). Nevertheless, preservation of testicular morphology should not necessarily be interpreted as evidence of normal male reproductive function. Fertility depends not only on sperm production but also on sperm maturation, transfer efficiency, mating behaviour, and overall physiological condition. Since these parameters were not directly assessed, the possibility remains that chronic PIB-MNPs exposure impaired male reproductive performance through functional rather than structural mechanisms. Similar observations have been reported in environmental toxicology studies where behavioural deficits contribute substantially to reduced reproductive success despite minimal histological alterations in reproductive tissues.

A notable observation in our molecular and biochemical analyses is that the sodium dodecyl sulfate (SDS) vehicle control exhibited a detectable upregulation of *Stat92E* expression (∼2.05-fold) and a modest alteration in acetylcholinesterase (AChE) activity, yet produced no observable impairment in systemic phenotypic endpoints such as gut barrier permeability (Smurf assay), brain vacuolization, negative geotaxis, or overall survival. This divergence highlights the fundamental difference in sensitivity thresholds between sub-cellular homeostatic responses and macroscopic tissue pathology. As an ionic detergent, low-concentration SDS acts as a mild chemical surfactant that induces transient cell-membrane irritation along the midgut epithelium. In response, *Drosophila* rapidly mobilizes the JAK/STAT signaling cascade, driving *Stat92E* transcription as an adaptive, compensatory repair mechanism that preserves epithelial barrier integrity and prevents physical dye leakage or systemic morbidity. Similarly, detergent interactions with membrane-bound AChE during *in vitro* assays can subtly alter enzyme conformation or active-site kinetics without breaching the critical neurotoxic threshold necessary to induce synaptic failure, neurodegeneration, or locomotor decline. By contrast, exposure to PIB-MNPs combines this baseline surfactant presence with insoluble, sharp micro/nanoplastic flakes (<2–10 µm) that cause persistent mechanical micro-abrasion of the peritrophic matrix, chronic oxidative stress, and structural barrier breakdown. Thus, while SDS alone triggers an early, protective cellular stress response that successfully maintains physiological homeostasis, the addition of physical micro/nanoplastic particles overwhelms these compensatory repair pathways to drive systemic pathological and phenotypic decline.

## Conclusion

In conclusion, exposure to PIB-MNPs exerts multisystem toxicity in *D. melanogaster*, impairing survival, locomotor function, neurophysiology, reproduction, and gut integrity while activating inflammatory responses. The observed sex-specific susceptibility and non-monotonic dose–response patterns highlight the complexity of PIB-MNPs’ biological interactions and underscore the need for further investigation into their environmental and health risks. Thus, findings from our study indicate that prolonged PIB-MNPs exposure induces progressive neurotoxicity in *Drosophila melanogaster*.

## Acknowledgement

We appreciate Pratyusha Banik of Vellore Institute of Technology for her assistance with PIB-MNPs synthesis and the Advanced Facility for Microscopy and Microanalysis (AFMM) for providing appropriate facilities for this study.

## Authors Contribution

D.M.B., A.T.A., and R.S. conceived and performed the experiments, analyzed the data, drafted the original manuscript, and edited the text. U.N. secured funding, supervised and managed the project, provided resources, and reviewed and finalized the manuscript. All authors have read and agreed to the published version of the manuscript.

## Use of artificial intelligence tools

Claude AI, a large language model, was used to assist with revising the manuscript for clarity, logical flow, and readability. All experimental work, data analysis, interpretation of results, and scientific conclusions are entirely the work of the authors. No artificial intelligence tools were used in data collection, analysis, figure generation, or code writing.

## Ethical Approval

This study was carried out following the guidelines on care and use of laboratory animals of the Institutional Animal Ethics Committee (IAEC) at Indian Institute of Science (IISc), Bangalore.

## Consent to participate

Not applicable

## Consent to publish

Not applicable

## Data Availability

Data will be made available on request.

## Supplementary data

**Supplementary Figure 1:**
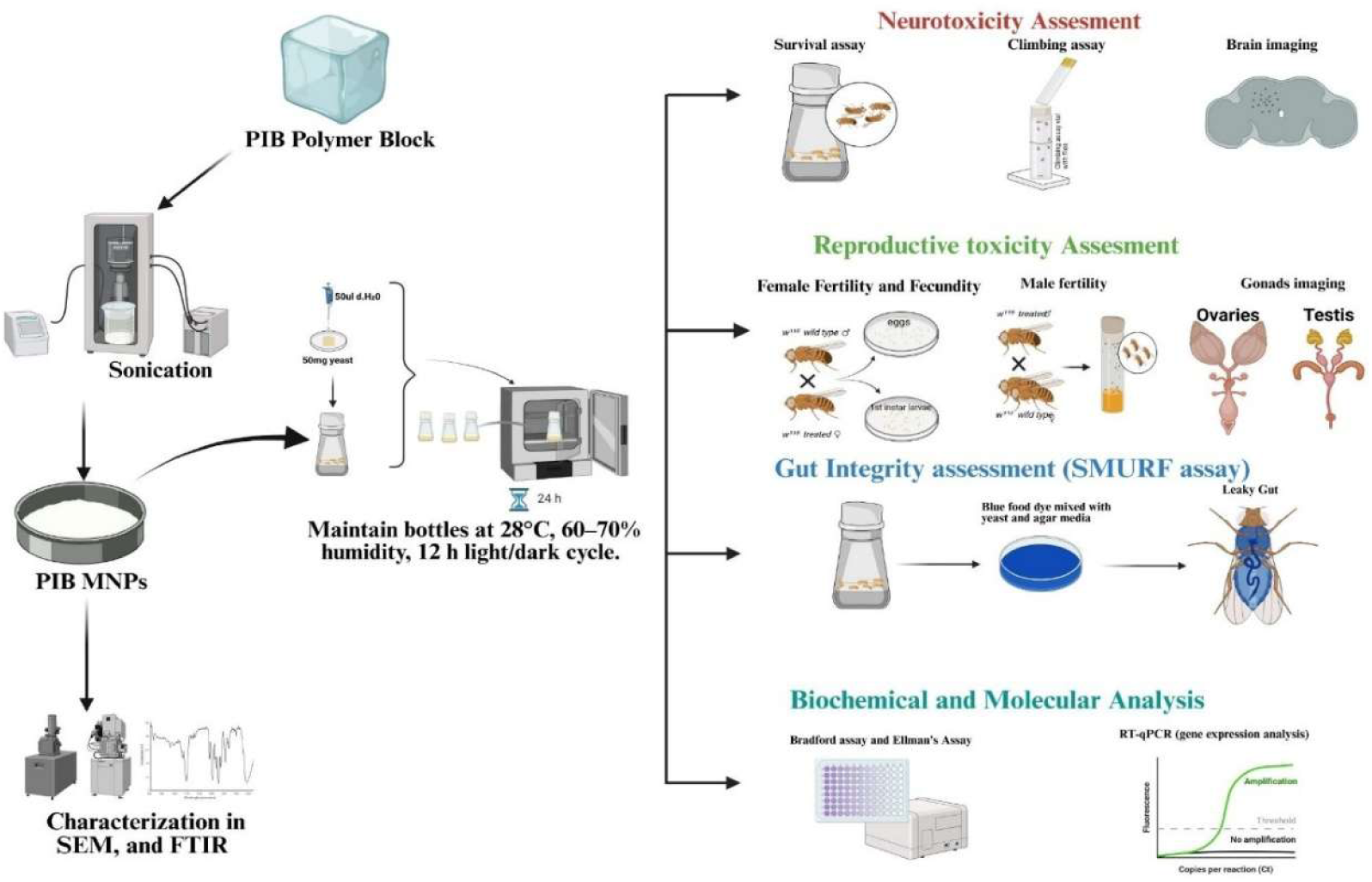
Schematic representation of the experimental workflow for evaluating the toxicity and biological effects of polyisobutylene (PIB) polymer nanoparticles (PIB-MNPs) in *Drosophila melanogaster*. PIB polymer blocks were sonicated to produce PIB MNPs, which were characterized using scanning electron microscopy (SEM) and Fourier-transform infrared spectroscopy (FTIR). Flies were maintained under controlled laboratory conditions (28°C, 60–70% humidity, 12 h light/dark cycle) with yeast-supplemented media. Toxicity endpoints included neurotoxicity (survival, climbing, and brain imaging assays), reproductive toxicity (fertility and fecundity assessments, gonad imaging), and gut integrity (SMURF assay using blue dye to detect intestinal leakage), all of which were affected. Biochemical and molecular analyses comprised Bradford and Ellman’s assays and RT-qPCR for gene expression profiling, which also confirmed the toxicity of PIB-MNPs. This integrated design provides a comprehensive approach to assess PIB nanoparticle-induced physiological and molecular alterations in a model organism.

**Supplementary Figure 2:**
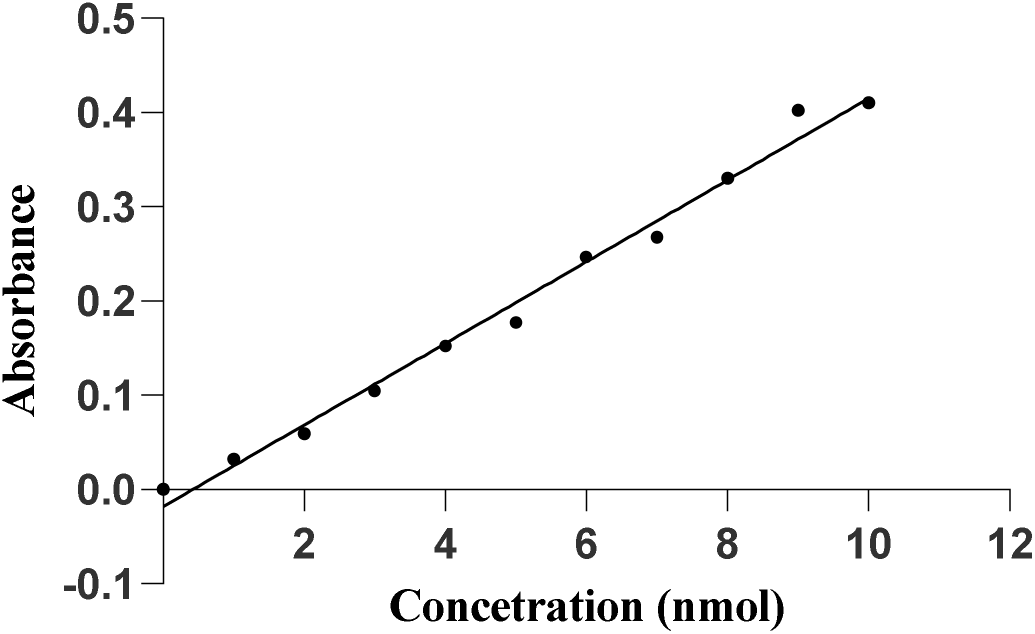
Glutathione (Ellman’s) Standard Graph

**Supplementary Figure 3:**
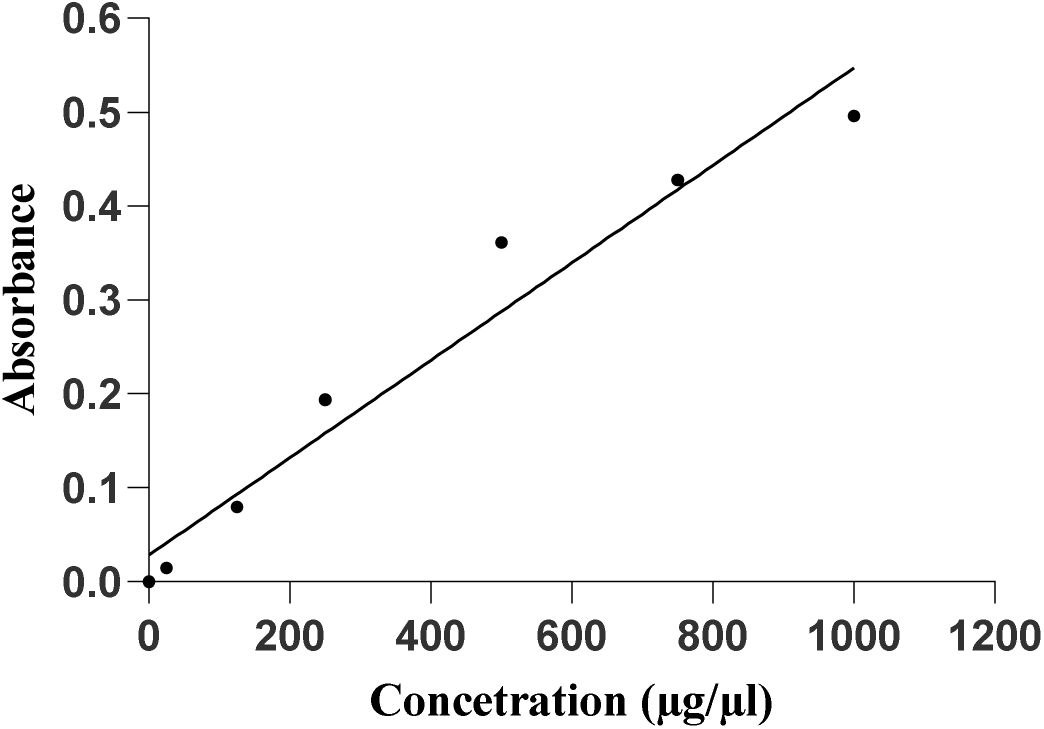
BSA (Bradford) Standard Graph

**Supplementary Figure 4:**
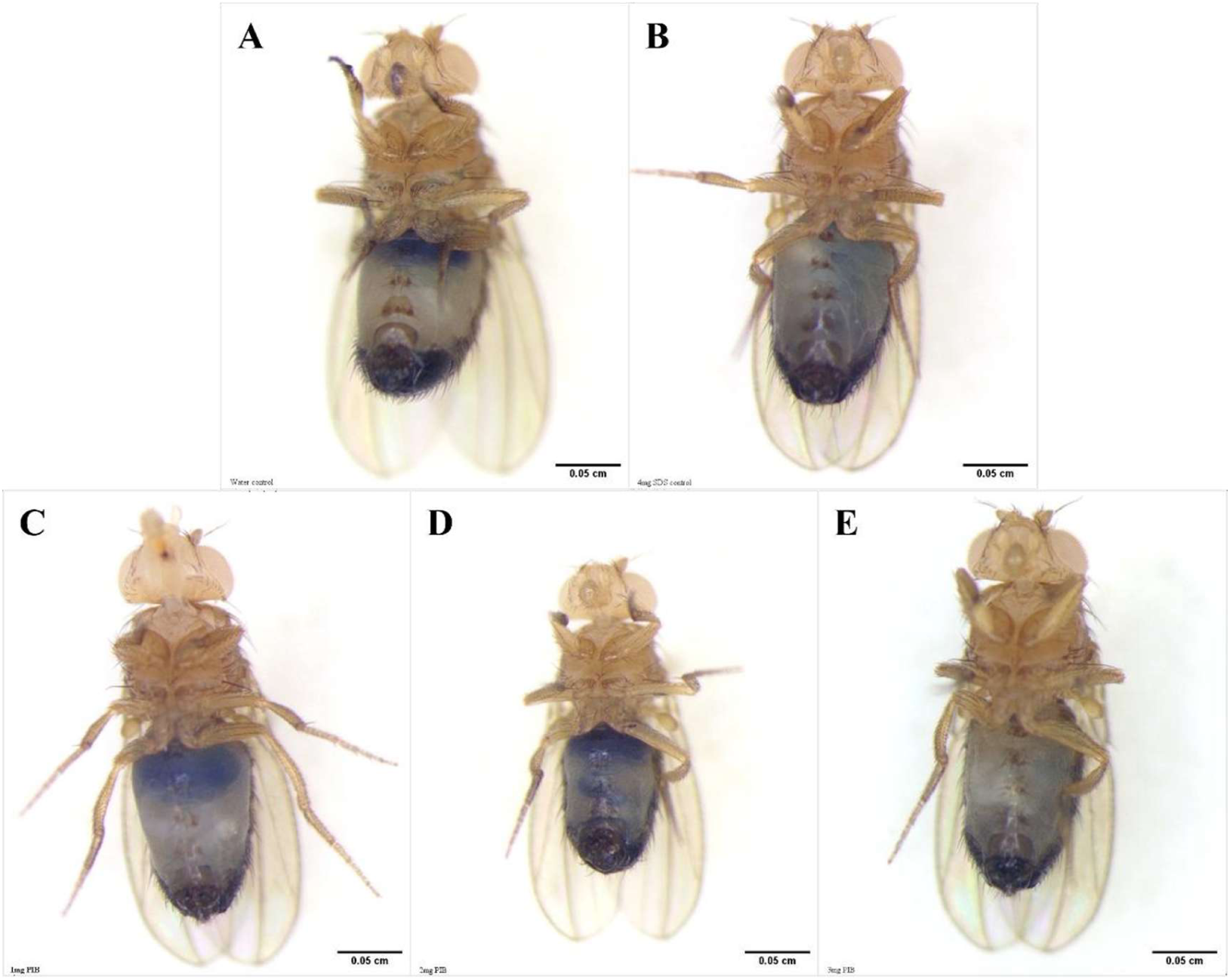
Representative stereomicroscope images of *D. melanogaster* following Smurf assay on day 21 of PIB-MNP exposure. Flies were fed blue dye incorporated into the yeast paste from day 5 until day 21 of the experimental period and examined under an Olympus SZX12 stereomicroscope (scale bar = 0.05 cm). Representative images from **(A)** water control, **(B)** SDS control, **(C)** 1 mg PIB-MNPs, **(D)** 2 mg PIB-MNPs, and **(E)** 3 mg PIB-MNPs treatment groups are shown.

**Supplementary Table 1:**
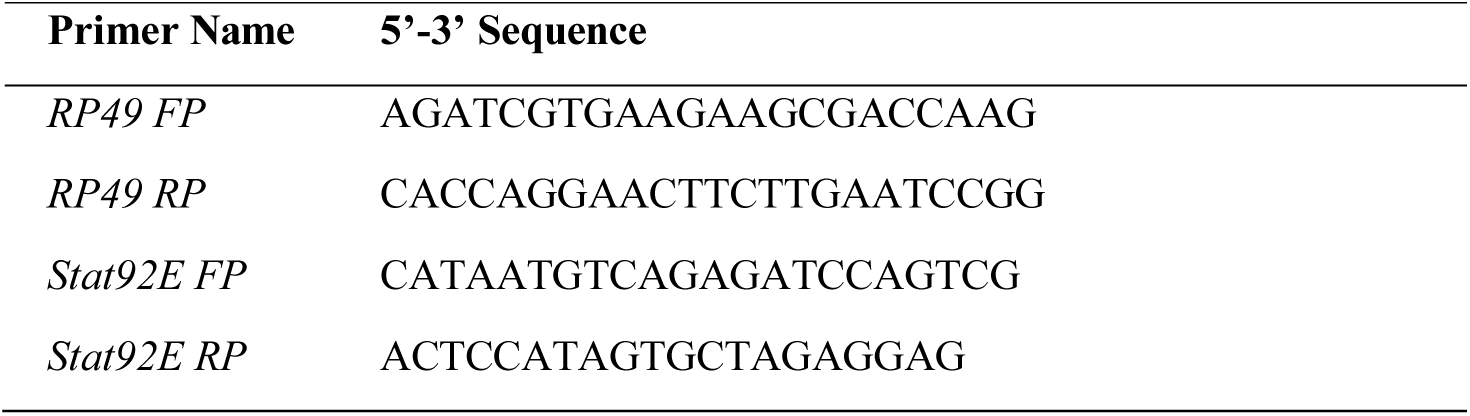
Primers used for RT-qPCR analysis.

